# Disrupted trans-placental thyroid hormone transport in a human model for MCT8 deficiency

**DOI:** 10.1101/2023.01.19.524727

**Authors:** Zhongli Chen, Selmar Leeuwenburgh, Wouter F. Zijderveld, Michelle Broekhuizen, Lunbo Tan, Rugina I. Neuman, Rutchanna M.S. Jongejan, Yolanda B. de Rijke, Irwin K.M. Reiss, A.H. Jan Danser, Robin P. Peeters, Marcel E. Meima, W. Edward Visser

## Abstract

Maternal-to-fetal transfer of the thyroid hormone T4 is essential for prenatal neurodevelopment, but the transporter facilitating trans-placental T4 transport is unknown. Mutations in the thyroid hormone transporter MCT8 cause a neurodevelopmental and metabolic disorder which key clinical features can be ameliorated by the T3 analogue TRIAC. The placenta has defective MCT8 if the fetus has MCT8 deficiency as the placenta is a fetal tissue. Should placental MCT8 be physiologically relevant, defective T4 transport across the placenta could represent a hitherto unrecognized mechanism underlying MCT8 deficiency. We investigated the importance of MCT8 and the trans-placental transport of TRIAC using an ex vivo human placental perfusion setup.

Our study (i) showed that MCT8 has a major role in maternal-to-fetal T4 transport, (ii) implies that disrupted placental transport of thyroid hormones could be the culprit in the early cascade of events in MCT8 deficiency and (iii) indicated that the T3 analogue TRIAC is efficiently transported across the placenta, independent of MCT8, holding potential in mothers carrying fetuses with MCT8 deficiency.

## Introduction

Thyroid hormones, the collective name for the prohormone T4 and the active hormone T3, are crucial for neurodevelopment. During prenatal neurodevelopment, maternal-to-fetal T4 transfer is critical, particularly during the first half of pregnancy with the fetal thyroid gland being immature (Patel *et al*, 2011). Therefore, maternal thyroid dysfunction negatively impacts brain structure and function in the offspring (Jansen *et al*, 2019; Korevaar *et al*, 2016; Li *et al*, 2010; Pop *et al*, 1999).

Intracellular bioavailability and transcellular transport of thyroid hormones are governed by plasma membrane transporters (Groeneweg *et al*, 2020). A key transporter is monocarboxylate transporter 8 (MCT8), which is expressed at the blood-brain barrier and in neural cells, and is crucial for transport of T3 and T4 (Friesema *et al*, 2003; Groeneweg *et al*., 2020). MCT8 deficiency, caused by mutations in MCT8, is a rare disorder consisting of severe intellectual and motor disability and abnormal thyroid function tests (Friesema *et al*, 2004). According to the current paradigm, the neurocognitive phenotype arises from impaired thyroid hormone entry into the brain (Lopez-Espindola *et al*, 2014; Vatine *et al*, 2017). With the blood-brain barrier being mature around 18 weeks, the placental barrier may be equally relevant for regulation of thyroid hormone bioavailability for the fetal brain. However, the transporter facilitating trans-placental thyroid hormone transport is unknown. Previous studies indicated MCT8 expression in the placenta (Chan *et al*, 2006). The placenta has defective MCT8 if the fetus has MCT8 deficiency as the placenta is a fetal tissue (Suryawanshi *et al*, 2018). Should placental MCT8 be physiologically relevant, defective thyroid hormone transport across the placenta (a fetal-derived barrier) could represent a hitherto unrecognized mechanism underlying MCT8 deficiency.

Direct postnatal administration of the T3 analogue TRIAC, which can bypass defective MCT8, normalized the brain phenotype in Mct8/Oatp1c1-knockout mice, the model resembling MCT8 deficiency (Kersseboom *et al*, 2014). Recently, we showed that TRIAC can ameliorate key clinical features of this disease (Groeneweg *et al*, 2019b). Therefore, it is paramount to assess its transport across the placenta for prenatal treatment in mothers carrying an MCT8 deficient fetus.

MCT8 is absent in commonly used human placental cell lines, largely limiting their role to study placental T4 transport (Chen *et al*, 2022b; Groeneweg *et al*, 2019a; Pan *et al*, 2021). Hence, we investigated the importance of MCT8 and the trans-placental transport of TRIAC in an ex vivo human placental model.

## Results

First, using western blot we confirmed expression of MCT8 in term placentas (**Figure EV1**). Previous studies reported that 10 µM silychristin completely inhibits MCT8-mediated T3 transport (Chen *et al*., 2022b; Johannes *et al*, 2016). Here, we determined that 10 µM silychristin also fully blocked MCT8-mediated T4 transport (**Figure EV2**).

Next, we investigated the role of MCT8 in trans-placental maternal-to-fetal transfer of T4 by adding 10 μM silychristin to the maternal reservoir in the presence of 100 nM T4 (**Figure 1A**). Pharmacological inhibition of MCT8 resulted in a ∼60% reduction of maternal-to-fetal T4 transfer after 3h-perfusion (4.2±1.2 nM fetal T4 in MCT8-inhibited placentas *versus* 10.6±0.6 nM fetal T4 in control placentas) (**Figure 1B**). As the deiodinase type 3 (D3) which converts intracellular T4 into the inactive metabolite reverse T3 (rT3) is highly active in human placenta (Koopdonk-Kool *et al*, 1996; Stulp *et al*, 1998), we also measured rT3 concentrations in the perfusates. Pharmacological inhibition of MCT8 resulted in a ∼42% reduction of rT3 concentration in the maternal circulation and a ∼80% reduction in the fetal circulation (**Figure EV3**). These findings indicate that MCT8 is responsible for a substantial amount of placental maternal-to-fetal T4 transfer.

**Figure 1.**
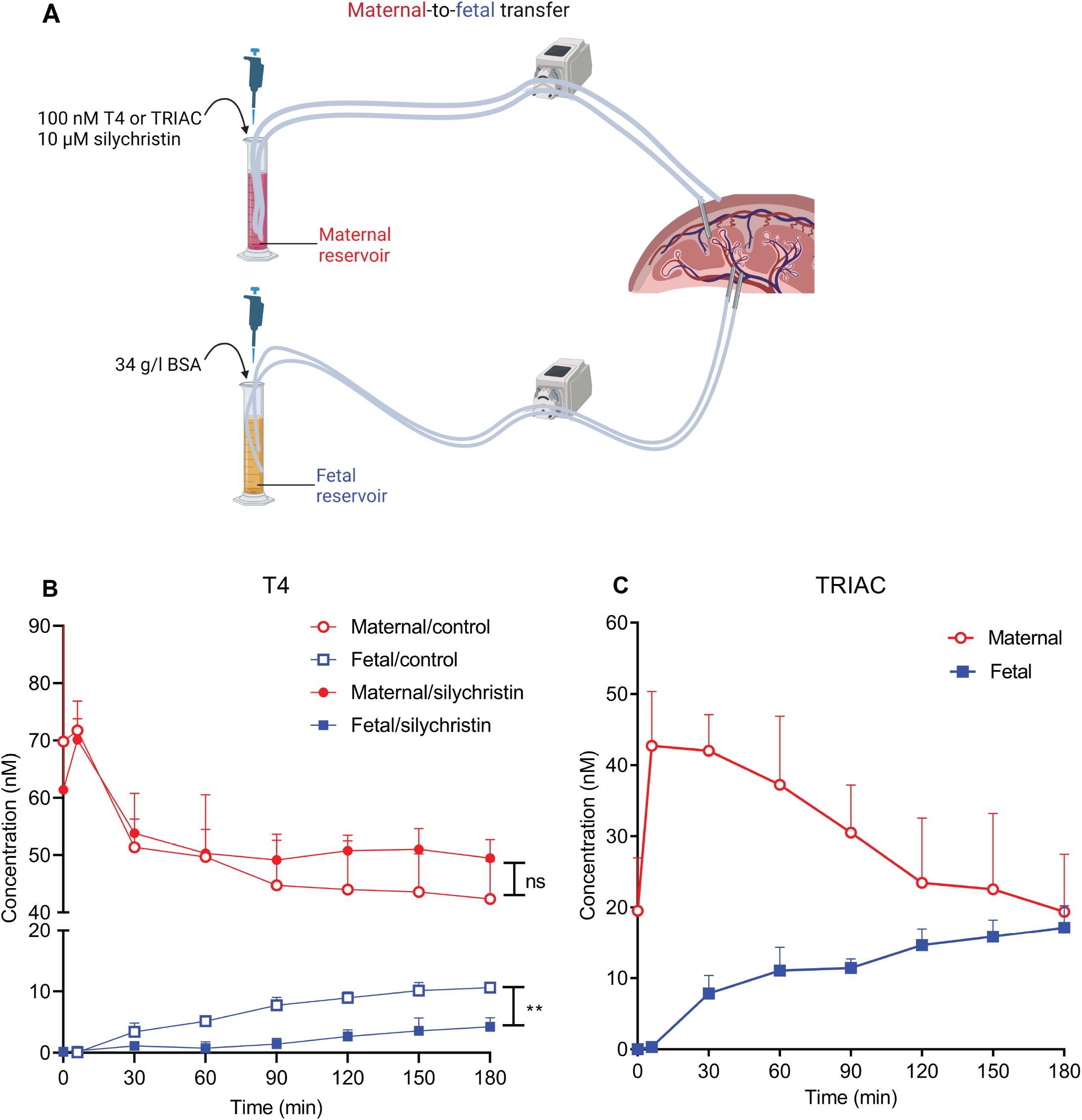
Maternal-to-fetal transfer of T4 or TRIAC in human placenta. (A) Schematic illustration of the *ex vivo* placental perfusion models. 100 nM T4 or TRIAC and 10 μM silychristin were added into the maternal reservoir and 34g/l BSA into the fetal reservoir. T4 (B) and TRIAC (C) concentrations from 3 perfusions for each condition. ns: not significant; **: p<0.01.

Finally, we utilized the pharmacological inhibition of MCT8 to mimic MCT8 deficiency in human placentas and tested trans-placental TRIAC transport (**Figure 1A**). TRIAC concentrations decreased from 42.7±6.2 nM to 19.4±6.6 nM in the maternal circulation and readily appeared in the fetal circulation (an increase from 0 nM to 17.1±2.5 nM in the fetal circulation after 3h-perfusion) (**Figure 1C**). These results show that TRIAC is efficiently transferred from the mother to the fetus in MCT8-deficient placentas.

## Discussion

Using an ex vivo human placental perfusion system, we showed that MCT8 has a major role in maternal-to-fetal transport of T4, potentially relevant for early fetal brain development. Therefore, impaired trans-placental T4 transport may be a new element in disturbing early brain development, even before maturation of the fetal BBB, in MCT8 deficiency. Indeed, bypassing the placental barrier with intra-amniotic LT4 administration in a pregnant woman carrying a fetus with MCT8 deficiency improved myelination and neurocognitive features compared to his untreated affected brother (Refetoff *et al*, 2021). Another implication of our findings is that compounds impairing MCT8 function during pregnancy (e.g. endocrine disrupting chemicals), may negatively impact on neurodevelopment in the offspring.

Following beneficial effects of TRIAC on metabolic outcomes, effects of early TRIAC intervention on brain outcomes in children with MCT8 deficiency are currently investigated (NCT02396459). The present study showing efficient maternal-to-fetal TRIAC transport provides preclinical support for future clinical studies of prenatal TRIAC treatment.

We acknowledge several limitations. First, we used human term placentas, while maternal-to-fetal T4 transport may be more relevant in early pregnancy. However, MCT8 is consistently expressed in placenta throughout pregnancy (Chan *et al*., 2006), favoring a role of MCT8 at all stages of pregnancy. Second, the inability to use binding proteins (e.g. bovine serum albumin (BSA)) at the maternal side, intrinsic to our model (Chen *et al*, 2022a), resulted in super-physiological free fractions of T4 and TRIAC in the maternal circulation.

Our study (i) showed that MCT8 has a major role in maternal-to-fetal T4 transport implying that compounds (e.g. endocrine disrupting chemicals) inhibiting MCT8 could also affect fetal T4 availability, (ii) implied that disrupted placental transport of thyroid hormones could be the culprit in the early cascade of events in MCT8 deficiency and (iii) indicated that the T3 analogue TRIAC is efficiently transported across the placenta, independent of MCT8, holding potential in mothers carrying fetuses with MCT8 deficiency.

## Materials and methods

### Reagents

T4, TRIAC, silychristin and bovine serum albumin (BSA) were purchased from Sigma-Aldrich (Zwijndrecht, the Netherlands). T4 and TRIAC were dissolved in 0.1N NaOH and silychristin in dimethyl sulfoxide (DMSO).

### Patients and placentas

The study received exemption for approval from the local institutional Medical Ethics Committee according to the Dutch Medical Research with Human Subjects Law (MEC-2017-418). All patients gave written consent before donating their placentas. Randomly selected placentas of uncomplicated singleton pregnancies were collected immediately after delivery (via cesarean section) at Erasmus University Medical Center, Rotterdam, the Netherlands. Retained placentas and placentas with maternal viral infections (HIV, hepatitis B, Zika) or fetal congenital abnormalities on ultrasound or maternal diabetes were excluded (Chen *et al*., 2022a; Hitzerd *et al*, 2019).

### Placental perfusion experiments

The perfusion model was set up as described before (Chen *et al*., 2022a; Hitzerd *et al*., 2019). Perfusion buffer consisted of Krebs-Henseleit buffer (118 mM NaCl, 4.7 mM KCl, 2.5 mM CaCl_2_, 1.2 mM MgSO_4_, 1.2 mM KH_2_PO_4_, 25 mM NaHCO_3_ and 8.3 mM glucose), supplemented with 5000 IU (0.5 ml/l) heparin LEO and aerated with 95% O2/5% CO2). Briefly, cotyledons from human term placentas were cannulated within 30 min after delivery of the placentas. The fetal and maternal circulations were established and kept running for ∼45 min as open circulations to wash out the blood of the placentas. The fetal flow rate was incrementally increased from 1 ml/min to 6 ml/min and maintained throughout the 3 hours’ experimental perfusion period. The maternal flow was kept at 12 ml/min. Both maternal and fetal circulations were closed prior to switching to 200 ml fresh perfusion buffer containing the substances described below. Antipyrine (Sigma) (final concentration 110 mg/ml) was used as a positive marker for the sufficient overlap between the maternal and fetal circulations as it is able to diffuse across the placental barrier. Fluorescein isothiocyanate–dextran (FITC-dextran) (Sigma, molecular weight 40 kDa) (final concentration 39.5 mg/ml) was used as a marker of integrity of the capillary bed (Chen *et al*., 2022a; Hume *et al*, 2004). Antipyrine was added to the maternal reservoir and FITC-dextran to the fetal reservoir.

To study T4 or TRIAC transfer from the maternal to fetal circulation, 200 μl of 100 μM T4 or TRIAC (final concentration 100 nM) and 200 μl of 10 mM silychristin (final concentration 10 μM) were added to 200 ml perfusion buffer in the maternal reservoir (**Figure 1A**). 6.8 g BSA (final concentration 34 g/l) was added to the fetal reservoir (200 ml). Antifoaming A concentrate was applied to the top edge of the cylinders to prevent excessive foaming. Placentas were perfused for 3 hours and 1 ml (T4 transfer experiments) or 2 ml (TRIAC transfer experiments) samples were collected from the reservoirs into blood collection tubes (BD vacutainer) at the time points indicated in the figures. The samples were centrifuged at 3000 rpm for 10 min and the supernatants were collected as perfusates and stored at -20 °C.

The concentrations of antipyrine and FITC-dextran in the perfusates were measured as described previously (Chen *et al*., 2022a; Hitzerd *et al*., 2019). The perfusion experiments were excluded from further analysis when the foetal-to-maternal (F/M) ratio of antipyrine was <0.75 at *t* = 180 min (indicating insufficient overlap between maternal and fetal circulation), or the maternal-to-foetal (M/F) ratio of FITC-dextran was >0.03 at *t* = 180 min (indicating compromised intactness of the placental barrier) (Chen *et al*., 2022a; Hume *et al*., 2004).

The data of the control group of T4 perfusion experiments were from our previous study (Chen *et al*., 2022a) which used the same procedure of placenta selection and perfusion protocol.

### Radioimmunoassays

T4 and reverse T3 (rT3) concentrations in perfusates were measured using radioimmunoassays as described previously (Chen *et al*., 2022a; Hume *et al*., 2004).

### LC-MS/MS measurement

TRIAC concentrations in perfusates were measured by liquid chromatography–mass spectrometry (LC-MS)/MS as described previously with some minor changes (Jongejan *et al*, 2020). The maternal perfusates were diluted 10 times with perfusion buffer and the fetal perfusates were diluted twice prior to LC-MS/MS measurement to avoid saturation. 500 µl of these diluted samples were used for measurement and the calibration curve was established in methanol.

### Statistics

Data are presented as mean ± SD of 3 placentas. GraphPad Prism 8.4.0 (GraphPad, La Jolla, CA) was used for data analysis and unpaired two-tailed t-test was used to compare T4 or rT3 concentrations in the absence or presence of silychristin at t=180 min. A p value <0.05 was considered significant.

## Acknowledgements

We thank all participating women for donating their placentas. We thank Sam Schoenmakers and his team for facilitating placenta logistics and Barbara M. van der Linden, Lotte W. Voskamp, Linda AI-Hassany and Ilse R. de Goede for obtaining informed consent. We thank the department of pharmacy for measuring antipyrine. This research is funded by the EU Horizon 2020 programme, ATHENA project, grant number 825161, which is gratefully acknowledged. This publication reflects only the authors ‘ view, and the European Commission is not responsible for any use that may be made of the information it contains.

## Author contributions

ZC, WEV, MEM, RPP: study design. ZC, MB, LT, RIN: perfusion experiments. ZC, LS: radioimmuno-assays. WFZ: LC-MS/MS measurements. ZC, MEM, WEV: data analysis and interpretation, writing of manuscript. All: critically review and approval of the manuscript.

## The Paper Explained Problem

Thyroid hormones (prohormone T4 and bioactive hormone T3) are essential for neurodevelopment. During prenatal neurodevelopment, maternal-to-fetal transfer of T4 is critical, particularly during the first half of pregnancy when the fetal thyroid gland is immature. Transcellular transport is governed by plasma membrane transporters, but the transporter facilitating trans-placental thyroid hormone transport is unknown. With the blood-brain barrier being mature around 18 weeks, the placental barrier may be equally relevant for regulation of thyroid hormone bioavailability for the fetal brain. Mutations in the thyroid hormone transporter MCT8 cause a neurodevelopmental and metabolic disorder which key clinical features can be ameliorated by the T3 analogue TRIAC. Should placental MCT8 be physiologically relevant, defective T4 transport across the placenta (a fetal-derived barrier) could represent a hitherto unrecognized mechanism underlying MCT8 deficiency. The T3 analogue TRIAC, which can bypass defective MCT8, can ameliorate key clinical features of this disease. Therefore, it is paramount to assess its transport across the placenta for prenatal treatment in mothers carrying a fetus with MCT8 deficiency.

## Results

We investigated the importance of MCT8 and the trans-placental transport of T4 and TRIAC using an ex vivo human placental perfusion model, simulating normal and MCT8 deficient placentas. We showed that inhibition of MCT8 greatly reduced maternal-to-fetal T4 transfer. Moreover, TRIAC was efficiently transferred from the maternal to fetal circulation independent of MCT8.

## Impact

First, we identified MCT8 as a major contributor to T4 transport across the human placenta. This observation not only fills a gap in physiology, but could have substantial implications (e.g. if MCT8 is affected through endocrine disrupting chemicals). Second, we discovered a hitherto unrecognized mechanism underlying MCT8 deficiency, emphasizing the relevance of prenatal treatment. Third, efficient maternal-to-fetal TRIAC transport provides preclinical support for future clinical studies of prenatal TRIAC treatment.

## Expanded View Figures

**Figure EV1.**
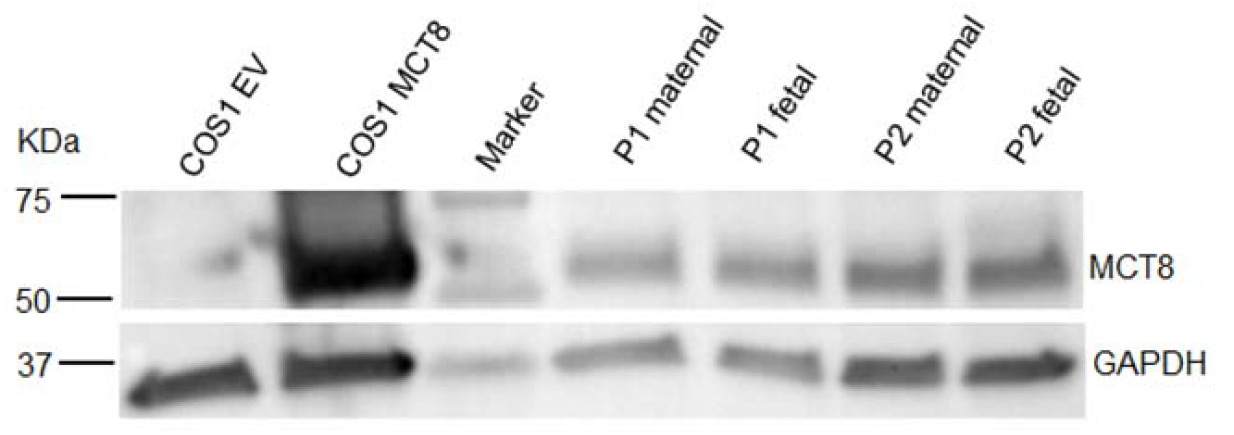
MCT8 protein expression in human term placenta. MCT8 was detected by western blot in the homogenates made from the biopsies that were collected from the maternal and fetal sides of 2 human term placentas (P1 and P2). Empty vector transfected COS1 cell lysate was used as a negative control and MCT8 transfected COS1 cell lysate as a positive control. Glyceraldehyde 3-phosphate dehydrogenase (GAPDH) was used as a loading control.

**Figure EV2.**
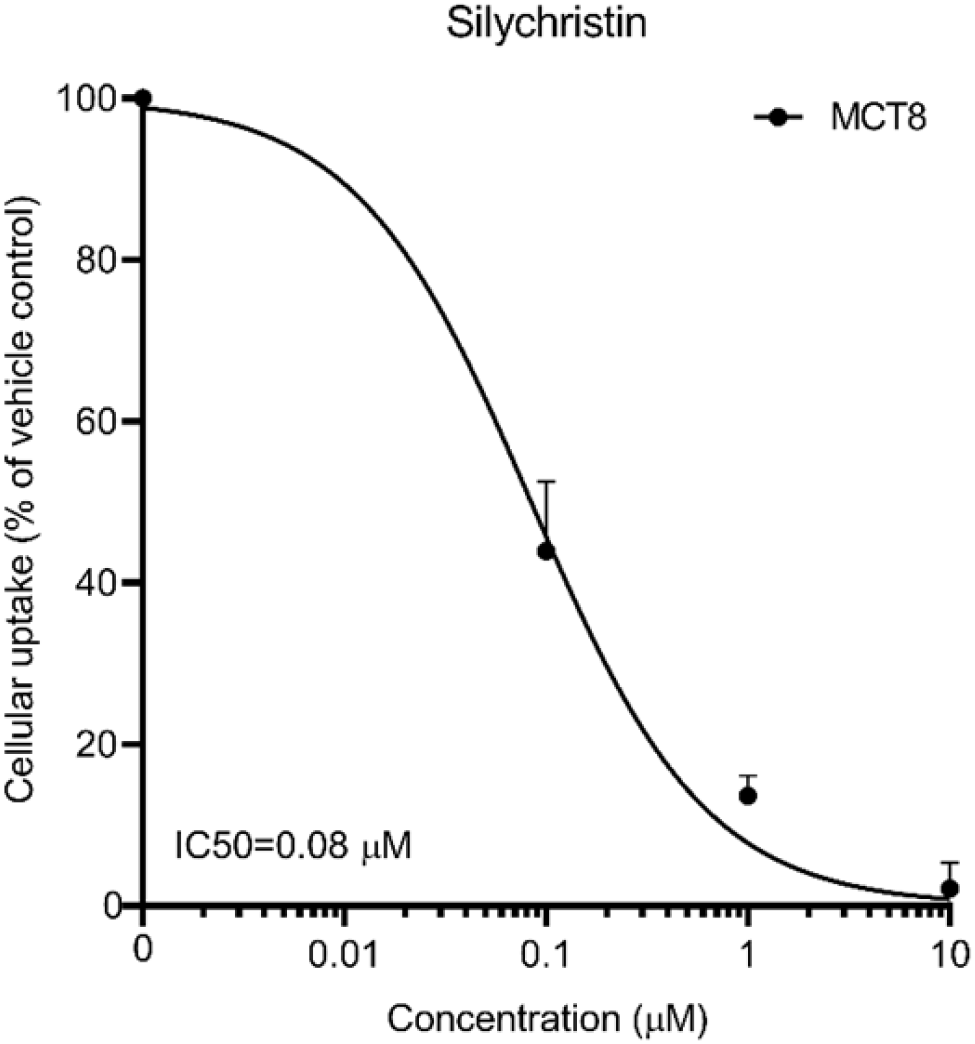
The efficacy of silychristin for T4 uptake in MCT8 expressed COS1 cells. COS1 cells were transfected with MCT8 or empty vector (EV) together with the intracellular thyroid hormone binding protein µ-crystallin (CRYM). Uptake assays were performed in DPBS/0.1%glucose with 1 nM ^125^I-T4 (50,000 counts per min) and incubated for 30 min. Uptake levels were corrected for background uptake in EV transfected control cells, incubated under the same condition and presented as a percentage. Resulting uptake levels were presented relatively to the uptake levels observed in presence of vehicle control. Data are presented as mean ± SD of 3 experiments. Non-linear regression (curve fitting) with the setting of “log (inhibitor) vs. normalized response” was used for the half-maximal inhibitory concentration (IC50) calculation (Chen *et al*., 2022b).

**Figure EV3.**
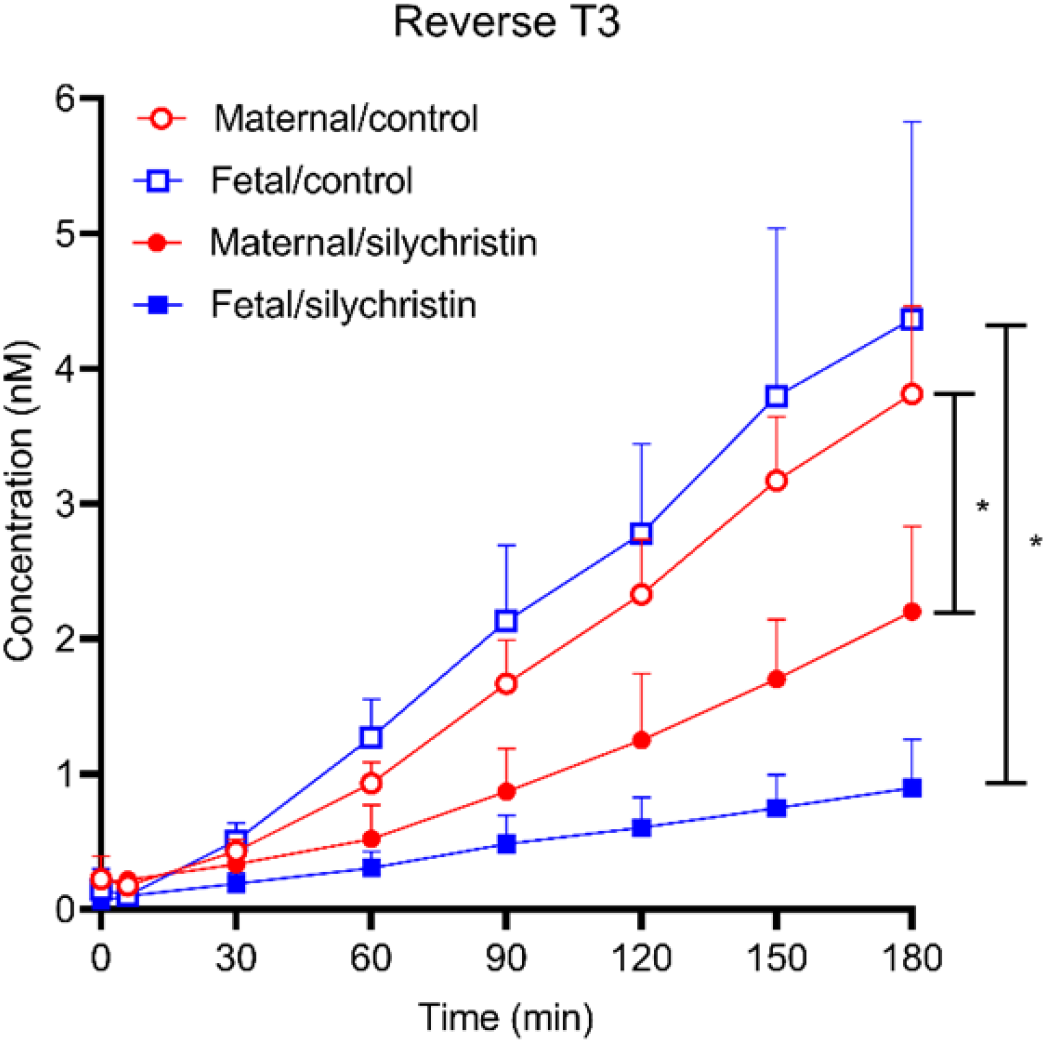
Reverse T3 (rT3) concentrations in maternal-to-fetal T4 perfusions in the absence or presence of silychristin. Samples were collected from the maternal and fetal circulations and rT3 concentrations were measured by radioimmunoassay. Data are presented as mean ± SD of 3 placentas and unpaired two-tailed t-test was used for statistical analysis to compare rT3 concentrations at t=180 min. * p<0.05.

